# Fractionating distraction: how past- and future-relevant distractors influence integrated decisions

**DOI:** 10.1101/2022.07.18.500552

**Authors:** Lydia Barnes, Dragan Rangelov, Jason B. Mattingley, Alexandra Woolgar

## Abstract

Many everyday tasks require us to integrate information from multiple steps to make a decision. Dominant accounts of flexible cognition suggest that we are able to navigate such complex tasks by attending to each step in turn, yet few studies measure how we direct our attention to immediate and future task steps. Here, we used a two-step task to test whether participants are sensitive to information that is currently irrelevant, but will be relevant in a future task step. Participants viewed two displays in sequence, each containing two superimposed moving dot clouds of different colours. Participants attended to one cued target colour in each display and reported the average direction of the two target dot clouds. In a subset of trials, we presented a “decoy” distractor: the second target colour appeared as the distractor in the first display. We regressed behavioural responses on the dot clouds’ motion directions to track how this future-relevant “decoy” distractor influenced participants’ reporting of the average target direction. We compared the influence of decoy distractors to never-relevant, recently relevant, and globally relevant distractor baselines. Across four experiments, we found that responses reflected what was immediately relevant, as well as the broader historical relevance of the distractors. However, relevance for a future task step did not reliably influence attention. We propose that attention in multi-step tasks is shaped by what has been relevant in the current setting, and by the immediate demands of each task step.

**Public Significance:** Our everyday functioning depends on our ability to piece together information to make coherent decisions. Understanding how we efficiently select and integrate goal-relevant information is critical if we wish to anticipate how decision-making can go wrong, whether because of fatigue, mental load, or illness. This study shows that decisions in multi-step tasks reflect two distinct processes: narrow focus on what is relevant in each step, alongside broader awareness of what has been relevant in this setting.

## Introduction

In everyday tasks, we often need to perform multiple, related, processing steps. For instance, to decide what fruit you would like to buy at the supermarket, you might find the fruit section, focus on the oranges, then hold the price of oranges in your mind while you look at the apples. Making an informed decision relies on having focused on each fruit in turn, maintained the relevant pricing information, and integrated it to select the best option.

This challenge of selecting and integrating multiple features is amplified by the fact that multiple goal-relevant features can be present at once. In the example of choosing fruit to buy, you may know from the start that you intend to choose either apples or oranges, so you need to extract some information about each. You then need to decide where to direct your focus at each moment so that information about the apples is not mixed up with information about the oranges.

Brain imaging, computational modelling, and behavioural research have highlighted that our capacity to self-impose periods of focus, or “attentional episodes”, around parts of a task, could be essential for fluid intelligence (Duncan, 2010, 2013; Duncan et al., 2017, 2020; Yang et al., 2019). This ability in turn powerfully predicts performance across a wide range of tasks (Pagani et al., 2017; Primi et al., 2010; Wray et al., 2020; Wrulich et al., 2014). Focusing exclusively on simple task parts could be especially important in novel or difficult tasks, where we cannot keep the whole task in mind (Duncan, 2013; Laird et al., 1986).

Conversely, approaching a sequential task by exclusively focusing on what is relevant at each moment could be burdensome. Neural network simulations show that engaging deeply with the current task can make it difficult to reconfigure the network and engage in a new task (Musslick et al., 2018). This is reinforced by a wealth of behavioural research on task switching, in which changing tasks typically produces slow and error-prone responses (Kray, 2006; Longman et al., 2017; Mayr & Keele, 2000; Meiran et al., 2000; Monsell, 2003; Rogers & Monsell, 1995). However, these findings only indirectly address the question of whether we proactively direct our attention to each part of a single task in turn, or to all relevant task features at once. As yet we do not have a reliable measure of how much we attend to non-immediate task parts.

In this study, we investigated people’s ability to form periods of focus around subsets of task-relevant information. We presented an integrated decision-making task (Rangelov & Mattingley, 2020), modified to test whether responses are biased by a target that appears outside the period in which it is relevant. Across four experiments, we found that attention was strongly biased by how often an item was relevant in general over the course of a few minutes. Once we accounted for this effect, however, attention was not reliably biased toward targets that appeared outside their period of relevance, that is, information that would be relevant in the subsequent epoch within the current trial. We propose that performance in simple sequential tasks is characterised by awareness of features that are relevant in the broader experimental context, together with sharp focus on each task part in turn.

## Experiment 1

### Methods

#### Participants

We modelled our task on Rangelov & Mattingley (2020), which showed that targets and distractors in multiple task epochs uniquely contributed to a behavioural response. We planned to explore whether distractor relevance modulated how targets and distractors contributed to behavioural responses. Thus, our key comparisons were between two distractor conditions, and we could not extract a likely effect size from Rangelov & Mattingley (2020). Instead, we designed this experiment to strongly contrast high and low distractor conditions, so that we could judge whether the task was sensitive to past- and future-relevant distractors before attempting to isolate different distractor influences. Our minimum effect of interest, then, was η2 = .06, meaning that distractor condition should explain more than 5% of the variance in target and distractor weights. In Cohen’s *d* terms, this means that distractor conditions should be at least half a standard deviation apart (*d*=.5).

We used G*Power (Faul et al., 2007) to calculate the appropriate sample size. We were specifically interested in whether future-relevant distractors (a) decreased the weighting of a first target and (b) increased the weighting of a first distractor on a behavioural response. Thus, we accounted for two one-sided, within-subjects t-tests, with alpha=.025 for each test (.05 total) and power of .7. For this first experiment, G*Power suggested that approximately 25 participants would be sufficient to detect our minimum effect of interest.

We recruited 25 participants (age=26.5±5.3 years, 17 female, 8 male) from the volunteer panel at the MRC Cognition and Brain Sciences Unit (MRC CBU). Participants could only access the study if they had previously reported that they were fluent in English, reported normal or corrected to normal visual acuity, reported normal colour vision, and were between the ages of 18 and 65. Two participants failed to meet the criteria for behavioural performance (see below) and were excluded, leaving 23 participants in the final sample (age = 26.39±4.97 years, 15 female, 8 male). Participants gave written informed consent before participating. All participants were given £6 per hour or pro rata, in minimum increments of £1.50, plus a small contribution towards travel costs (£2.50 or £3). The project was approved by the Psychology Research Ethics Committee at the University of Cambridge (PRE.2018.101).

#### Task

Participants completed a computer-based behavioural task (see Figure 1). Each trial began with a fixation cross for an intertrial interval between 500 and 1500 ms. A circular cue, presented for 500 ms at fixation, showed the target colours for the trial. The first target colour was shown in the left semicircle and the second target colour in the right.

**Figure 1.**
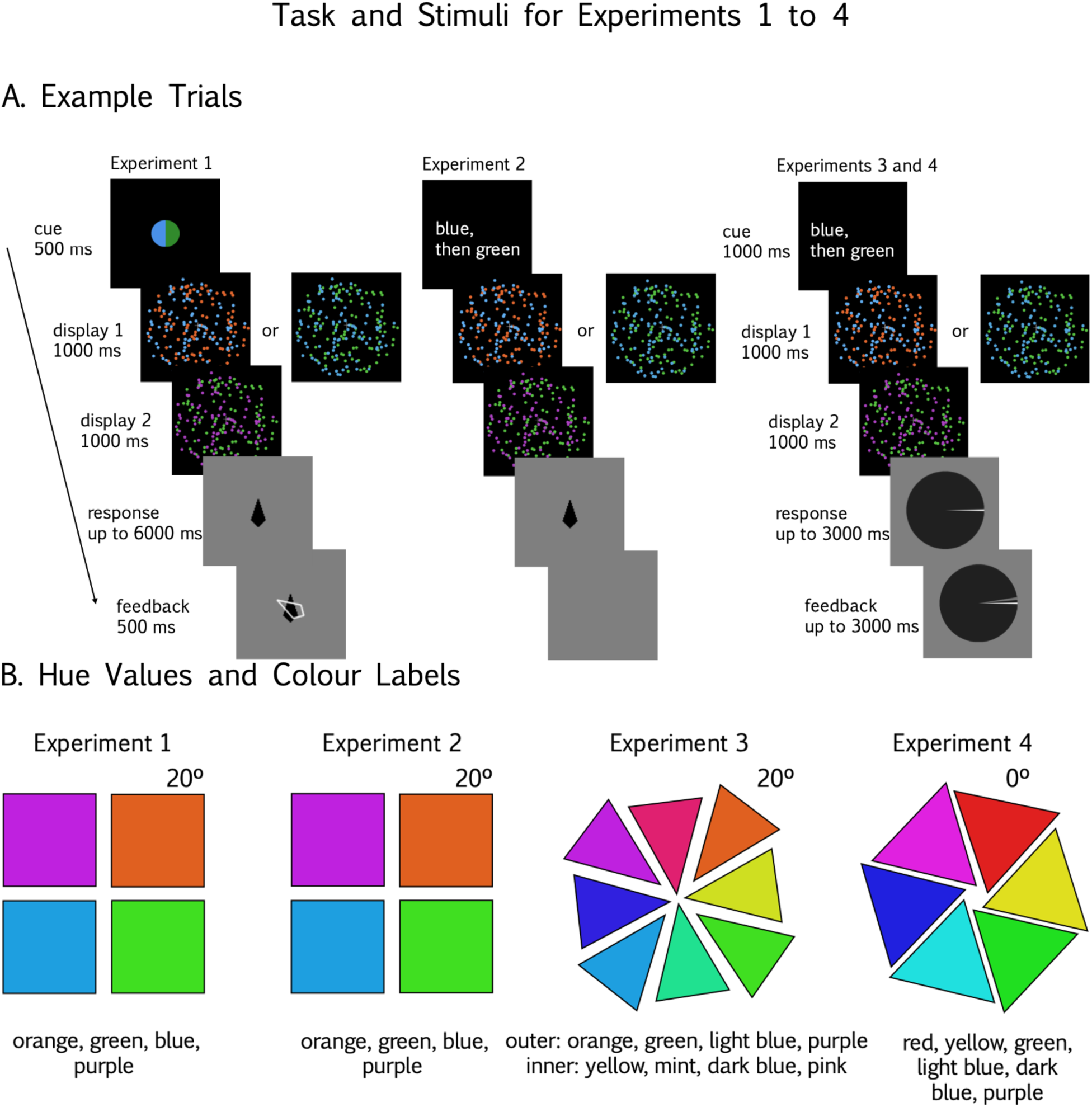
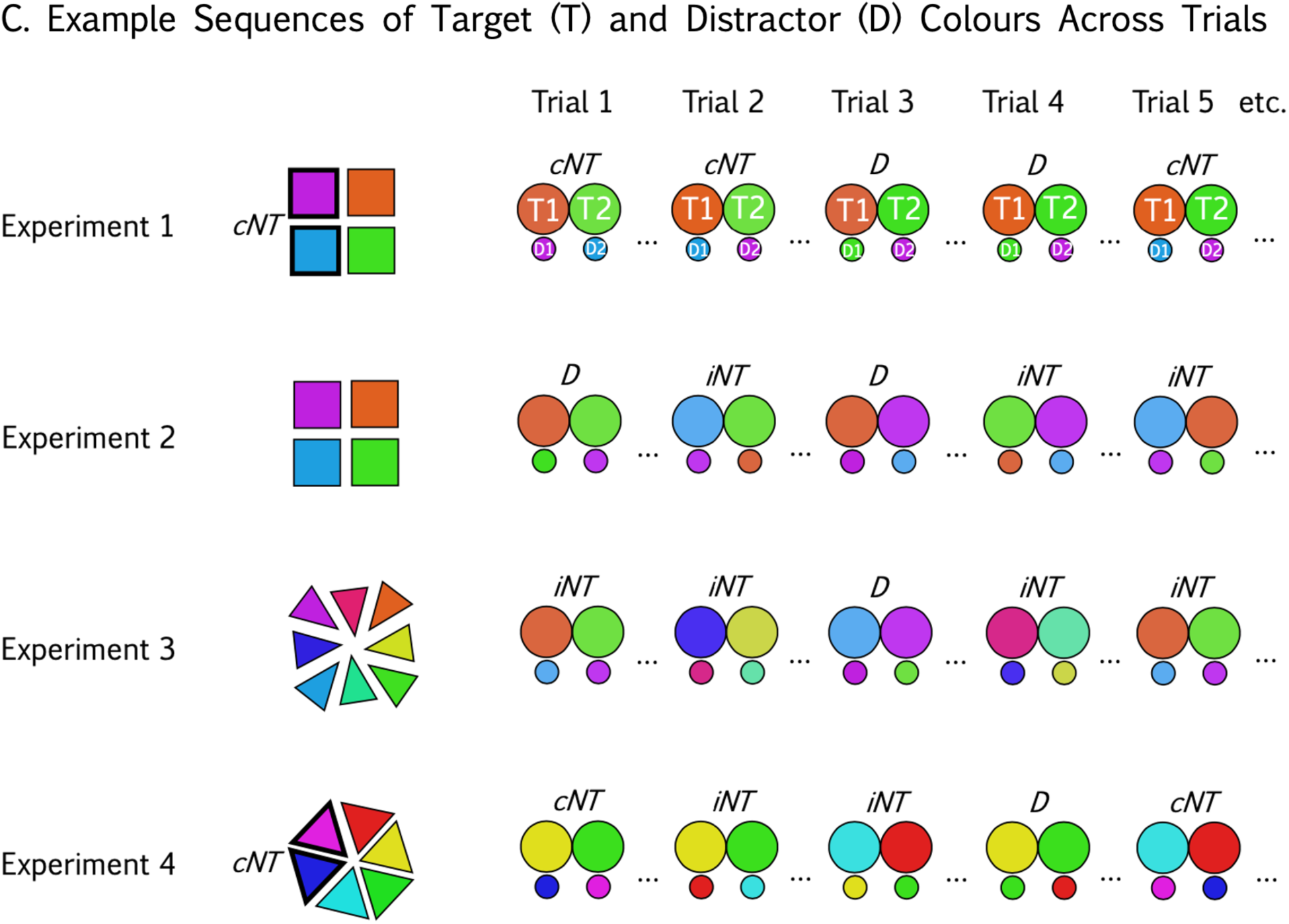
Schematics of trial structure and stimuli used in Experiments 1-4. (A) Example trials for Experiment 1 (left), Experiment 2 (middle), and Experiments 3 and 4 (right). The panel for Experiment 1 also shows example stimuli for baseline and decoy conditions. Experiments 1 and 2 had the same timings. (B) Colours for each experiment. The hue value is marked for the first colour in each set moving clockwise from 0° in HSL colour space. Subsequent hues in the set were equally spaced around the circle (i.e., 90° for Experiments 1 and 2, 45° for Experiment 3, and 60° for Experiment 4). Colour names below the colour sets were the labels used for written cues, where applicable. For Experiment 3, outer colours formed Set 1, and inner colours Set 2. (C) An example trial sequence for each experiment. Large circles represent Target 1 (left) and Target 2 (right) colours, with small circles representing distractors. Trials are labelled D for decoy, and cNT or iNT to indicate a baseline trial with never-relevant colours (“consistent non-targets”) or target colours (“inconsistent non-targets”) as distractors. Colours that are never relevant in the example sequence are labelled on the colour wheel where applicable (Experiments 1 and 4).

The cue was followed by two 1000 ms displays. Each display contained two moving dot clouds, one in a cued colour (the target). Participants remembered the motion direction of the target dot cloud on each display. After the dot displays disappeared, participants reported the average motion direction of the two target dot clouds by moving a black, obelisk-shaped response dial with a joystick. If participants did not respond within a 6000 ms time limit, the task continued automatically. After the response, or the maximum response time, a white feedback dial appeared imposed over the response dial, illustrating the correct response in a contrasting (black) colour for 500 ms.

We embedded two core conditions by manipulating the task-relevance of the un-cued dot clouds (the distractors). In the baseline condition, Distractor 1 or Distractor 2 were presented in a non-target colour that was irrelevant both to the current trial and to the task as a whole. That is, these baseline distractors were “consistent non-targets”. In the decoy condition, Distractor 1 was presented in the anticipated Target 2 colour (Figure 1A, left panel). For example, participants might see a colour cue with blue and green (as in Figure 1A). They would then see the first display containing two coloured dot clouds: the target dots in blue, and the distractor dots. Distractor dots could be in a consistent non-target colour (in this case, orange), or in the anticipated Target 2 colour, green. Participants would then see a second display with target dots in green, and distractor dots in a consistent non-target colour (in this case, purple).

Cues were blocked, and baseline and decoy trials were randomly mixed within each block with equal proportions of each condition. As we were primarily interested in cognitive processing of stimuli that would be relevant in the future, the decoy distractors, when present, always appeared in the first epoch.

#### Stimuli

We generated random dot stimuli in PsychoPy (Peirce, 2007; Peirce et al., 2019). Each dot cloud consisted of 80 dots, of which 40% moved in a coherent direction, well above the 10% coherence required for a typical adult human to detect motion. The remaining 60% followed a random walk. This ensured that participants could not rely on individual dots to indicate the motion direction, as any dot had a .4 probability of containing signal in one epoch and .16 probability of containing signal across two epochs. Dots moved at 19 degrees of visual angle per second. Each dot was randomly assigned a duration between 0 and 100 ms. When a dot reached its assigned duration, it was replaced by a new dot at a random location.

We selected a range of colours to distinguish superimposed dot clouds. All colours were set at 75% saturation and 50% luminance in HSL (hue, saturation, luminance) colour space. We selected four colours with hues 90° apart, starting from 20° (Figure 1, Panel B). We adjusted these colour values for colour distortion to ensure that saturation and brightness were matched on the lab computer monitors.

Motion directions were selected from 0 to 359° in steps of 1°. Target and distractor motion within an epoch (e.g., Target 1 and Distractor 1) as well as the two target motions between epochs (e.g., Target 1 and Target 2) could differ by between 30 and 150°. This ensured that the average of the target dot clouds’ motion direction gave an unambiguous answer (that is, target clouds were never separated by 180°).

Stimuli were presented on a 19-inch Dell 1908FP LCD monitor at 1280×1024 resolution and refresh rate of 60Hz. Stimuli subtended 15.94 degrees of visual angle at a viewing distance of 50 cm.

#### Procedure

Participants were seated comfortably at a computer monitor. All participants gave basic demographic information in a pre-experiment questionnaire (age, sex, vision). They then began training on the task. Training consisted of three blocks of 24 trials each. In the first block, participants saw only a single epoch of coloured dots and reported the target motion direction presented in that epoch while ignoring concurrently presented distractor motion. In the subsequent two blocks, they practiced the dual-epoch task. Cue durations began at 1000 ms in the first two blocks, reducing to 500 ms in the third training block, to ease participants into the task. The experimenter discussed errors with the participant between training blocks and explained any aspects that were unclear.

Following training, participants began the main session. They completed 12 blocks of 384 trials. Each participant was presented all possible cues. That is, each of the four colours was paired once with each other colour (orange with green, orange with blue, etc.) to form six cues. These cues were reversed (green with orange, blue with orange, etc.) to form six more cues and create 12 blocks. Block sequence was randomised within and between participants.

##### 4.2.1.5. Analyses

###### Data quality

Participants with incomplete data, or whose mean absolute error exceeded 45°, were excluded from the analysis.

###### Ordinary least-squares regression

For each participant, we regressed their responses on the true motion directions of the target and distractor dot clouds. Since motion directions were angles, and real-valued angles are circular (-π is the same as π), we represented angles for the predictors (design matrix, X) and responses (R) as complex values (Eq. 1). We then fit a linear model (Eq. 2), using ordinary least squares regression (Eq. 3) to estimate the weight vector 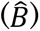 that optimally fit the four predictors to the responses.

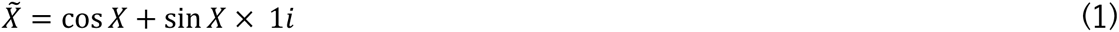

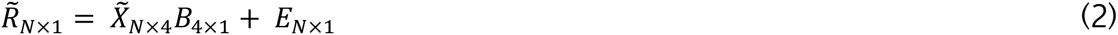

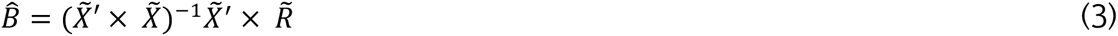

The weight matrix, like the predictors and responses, was complex valued. We took the absolute values of the weight vector (Eq. 4). These values represent an expansion factor for each predictor, so that a high weight for a given predictor represents a strong influence on the response. We refer to these absolute weights as “decision weights” because they reflect an estimate of how the target and distractor dot motions influenced participants’ choices at the response screen.

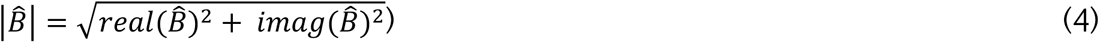

###### Permutation testing

Next, we obtained single-subject measures of attention to Target 1 and Target 2. Decision weights could not be negative, meaning that chance values could be greater than zero but not below. Rather than test the decision weights against a normal distribution, which assumes that chance values are distributed around the null value, we tested the decision weights against individual-specific, permutation-based null distributions. We took each participant’s predictor values and randomly permuted them, so that target and distractor dot directions for a given trial could be assigned to any trial. We calculated the decision weights as above for these permuted data, and repeated the process until we had decision weights for 10,000 permutations. This formed the empirical null distribution of decision weights for that participant. We then compared the decision weights for the correctly labelled (unpermuted) data to the null distribution. Decision weights that exceeded the 95^th^ percentile of the null distribution were considered unlikely under the null hypothesis, and reliably different from zero.

The main purpose of this analysis was to confirm that participants were doing the task as instructed, using the target decision weights as a secondary measure of accuracy. In particular, participants could respond somewhat accurately by attending to one of the two targets on each trial. A strategy of attending to either Target 1 or Target 2 would make it difficult for us to judge how a decoy distractor, presented in the Target 2 colour, influenced focus on Target 1. Thus, we planned to use these individual-level tests as an additional exclusion criterion for participants whose baseline target weights were not both reliably different to zero.

###### Group baseline analysis

Next, we compared baseline target and distractor weights to zero at the group level. As individual participants were excluded if their target weights did not both exceed zero, target weights should always exceed zero across the group. However, just as target decision weights provide information about the source of accuracy, distractor decision weights can provide information about the source of inaccuracy: in this case, whether errors were random, or reflected attentional capture by Distractor 1 or Distractor 2. Understanding this could provide context for comparing conditions, for which we expected distractor weights to play a role. Thus, we tested whether each decision weight was greater than zero in the baseline distractor condition. For each condition and dot cloud, we collated the individual-level null data and estimated a cumulative distribution for the null across the group. If the group’s average decision weight exceeded the null value whose probability was 5% in the null distribution, the decision weight could be considered reliably different to chance at the alpha = .05 level. To account for multiple comparisons, we applied a family-wise (Bonferroni) correction. We set alpha at .013 (0.05/4) for one-sided comparisons of each decision weight to zero.

###### Condition comparisons

Finally, we compared target and distractor weights between the conditions with a paired t-test on baseline and decoy conditions. If the future-relevance of an upcoming target affected the current attentional set, we expected that decision weights would be higher for Distractor 1 in the decoy condition relative to baseline. On the same assumption, we expected lower weights for Target 1 in the decoy condition relative to baseline. Consequently, we planned two one-sided t-tests, setting alpha at .025. For completeness, we analysed Target 2 and Distractor 2 weights in the same way, but do not interpret these data as we did not have targeted hypotheses for these comparisons.

### Results

#### Data quality exclusions

Two participants were excluded for high errors, leaving 23 participants (67.2% power to detect our effects of interest).

#### Baseline decision weights

All remaining individual subjects’ baseline target weights exceeded zero, compared with their individual permutation-based null distributions. Baseline target and distractor decision weights across the group were also reliably different from zero (all p<.001, alpha=.013).

#### Decoy effect on decision weights

We predicted that if momentary attentional focus was affected by the upcoming attentional set, Target 1 weight would decrease, and Distractor 1 weight would increase, in the presence of a ‘decoy’ distractor.

Indeed, Target 1 weights decreased (*d*=1.27), and Distractor 1 weights increased (*d*=-.85), when the distractor in Epoch 1 was the anticipated Target 2 colour, relative to a non-target baseline (Figure 2). That is, within a trial responses were less influenced by relevant information, and more influenced by irrelevant information, when the distracting information in Epoch 1 was relevant to the subsequent task in Epoch 2. We also observed a similar pattern for Target 2, whereby having just ignored a particular colour in the preceding epoch of the trial (decoy condition) reduced the weight assigned to that colour in the current epoch.

**Figure 2.**
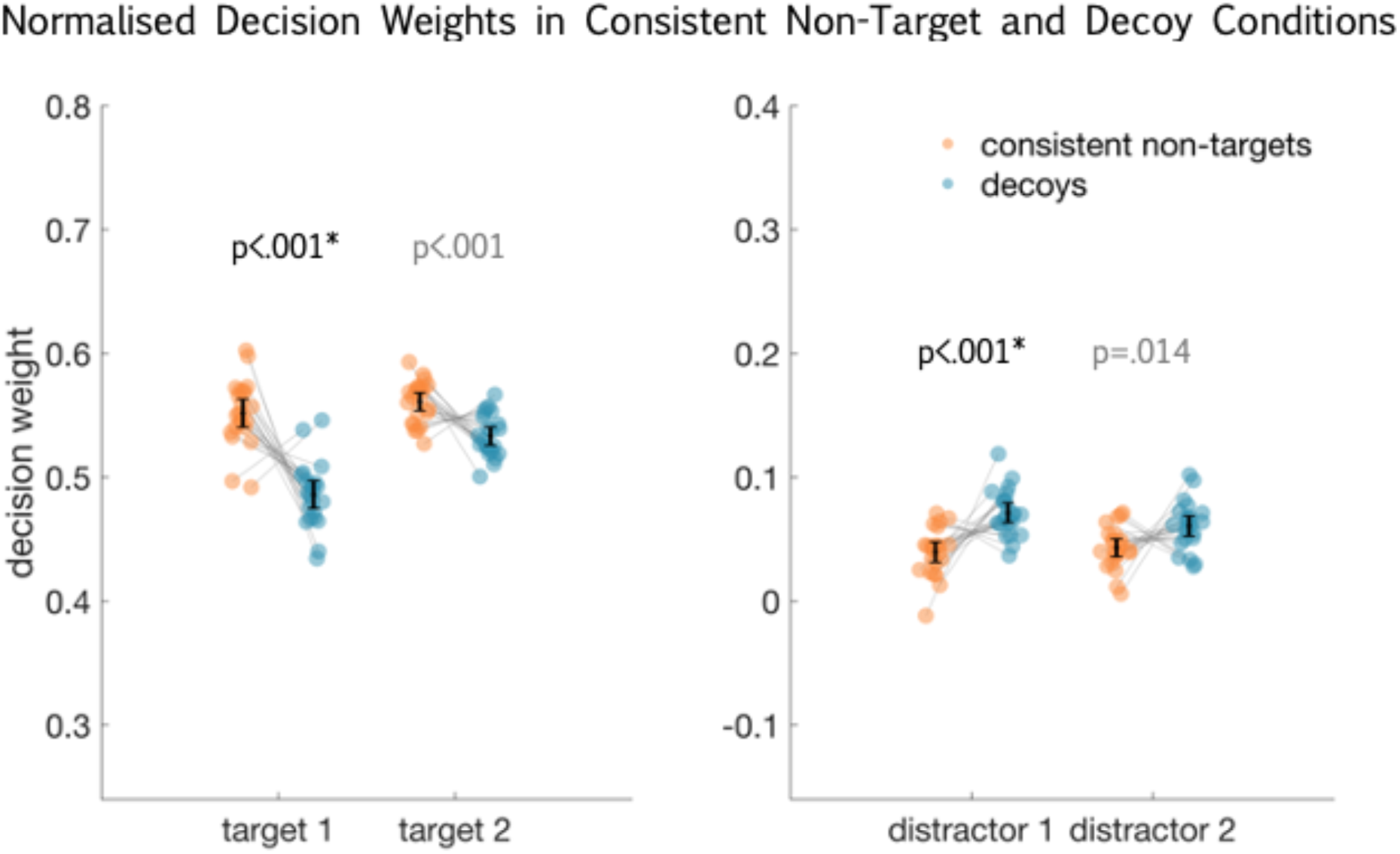
Decision weights for target and distractor dot clouds for Experiment 1, shown separately for consistent non-target baseline (orange) and decoy (blue) conditions. Decision weights are normalised for visualisation, and show within-subject effects centred on the group mean (Cousineau, 2005). Grey lines between conditions connect individual participants. Black bars indicate the 95% confidence interval. P-values are derived from paired t-tests and are uncorrected for multiple comparisons. Between-condition comparisons for Target 2 and Distractor 2 are in grey. Statistically significant condition differences in our planned comparisons are marked with an asterisk.

These findings are consistent with the idea that current attentional focus is influenced by the upcoming relevance of related information; that is, that attention is not allocated in strictly discrete temporal chunks. It appeared that participants would down-weight relevant information, and up-weight irrelevant information, when the irrelevant information was presented in the anticipated target colour of the subsequent epoch. This pattern also extended to a second task epoch, suggesting that suppressing a future-relevant distractor has negative consequences for attending to that stimulus soon after.

However, distraction by the decoy in this experiment could also be driven by factors besides its imminent relevance in the subsequent epoch. First, cues on each trial displayed the target colours on screen immediately before the trial, meaning that the decoy distractor was both future-relevant and recently seen. Distraction by a decoy because it was cued is difficult to disentangle from distraction due to its future relevance for the task, since its future relevance is by definition determined by the cue. However, the distinction is important if we want to argue that decoy effects here specifically reflect how we manage future-relevant information.

Second, the task used a single cue (for example, “attend to orange, then blue”) for a full block. This meant that the decoy was both the anticipated Target 2 colour, and the Target 2 colour that participants had responded to on the previous trial. It is possible, therefore, that higher Distractor 1 weights for decoys relative to baseline reflected effects of recent history rather than preparation for the immediate future. Finally, the baseline distractor colours were constant over each block, making them globally less relevant to the task than the colours used for targets and decoy distractors. To summarise, then, there are three possible explanations for the decoy effect in Experiment 1: sensory priming or carry over of attentional set from the previous trial (recent relevance), up-weighting colours that are globally relevant across the task (global relevance), or preparing to attend to the upcoming target (future relevance). We therefore carried out a series of further experiments to isolate the role of future relevance from these alternative explanations.

## Experiment 2

Although we saw the anticipated effect of decoys in Experiment 1, we could not disentangle the contribution of recent and global relevance from the decoy’s future relevance for the trial. We designed Experiment 2 to isolate the influence of future-relevance on attentional processing, by equating recent and global relevance between decoy and baseline conditions.

### Methods

#### Participants

We again conducted a power analysis to estimate an appropriate sample size. Based on Experiment 1, we could expect distractor conditions to substantially affect target and distractor weights. However, the purpose of the current experiment was to isolate one factor that could have contributed to the condition difference that we observed in Experiment 1. Thus, although effects in Experiment 1 were large, we wanted to increase our sensitivity to detect smaller effects. We set our minimum effect of interest at η2 = .03. That is, we were interested in detecting distractor effects that explained at least 3% of the variance in behavioural responses. This equates to a Cohen’s *d* of .35. As in Experiment 1, we estimated the sample size for two within-subjects t-tests, alpha = .025 per test, power = .7. G*Power suggested that approximately 50 participants would be sufficient to detect our minimum effect of interest. We recruited an independent sample of 48 participants (age=33.1±15.0 years, 30 female, 18 male) for this study, through the MRC Cognition and Brain Sciences Unit Volunteer Panel. Payment and recruitment were as in Experiment 1.

#### Task

As in Experiment 1, we presented distractors in a colour that was irrelevant for the trial (baseline) or in the anticipated target colour (decoy). All task and analysis features remained the same as in Experiment 1, except for the following changes, which we introduced to match the broader task-relevance of baseline and decoy distractor colours.

First, we presented colour cues in text (for example, the words “purple, then blue”; see Figure 1, Panel A, centre). We did this to ensure that participants actively constructed their attentional biases, and did not simply respond to targets and decoys because they had recently seen those colours in the cue.

Second, we reduced the opportunity for positive trial-to-trial carry-over of attention (recent relevance), and equalised it between decoy and baseline conditions. In Experiment 1, trials were blocked, so that a decoy distractor in the first epoch – the second target colour on the current trial – was also the second target colour from the previous trial. To minimise this, in Experiment 2 we randomly selected a cue on each trial, so that target colours varied within a block. Randomly assigning each colour to be a target or distractor on each trial meant that carry-over of attentional biases from trial-to-trial could be positive or negative. That is, carry-over could equally help or hinder attention to targets and decoys. More importantly, the opportunity for carry-over was now matched between baseline and decoy conditions. We reasoned that, if participants still upweighted the decoy, relative to the baseline, distractor, this could no longer be easily explained by the decoy’s recent history of relevance.

Third, we manipulated the global relevance of the baseline distractors across the task. In Experiment 1, baseline distractor colours were consistent across a block, and were never used as target colours (“consistent non-targets”). In this experiment, the random cue sequence meant that target colours on one trial could become baseline distractor colours on the subsequent trial (“inconsistent non-targets”). Thus, participants could no longer benefit from a tendency to up-weight globally relevant colours, as the same colour appeared equally as often as a distractor or target in the baseline condition.

### Results

#### Data quality exclusions

Eight participants were excluded for high errors, leaving 40 participants in the final sample (age = 32.38±15.37, 23 female, 17 male; 57.5% power to detect our effects of interest).

#### Baseline decision weights

All remaining subjects’ baseline target weights exceeded zero, compared with their individual permutation-based null distribution. Baseline target decision weights across the group were also reliably different from zero (p<.001). In contrast to Experiment 1, baseline distractor decision weights were no longer statistically greater than zero.

#### Decoy effect on decision weights

In the blocked design of Experiment 1, decoy distractors were more distracting than consistent non-target distractors. Here, we used a random trial sequence to ensure that baseline distractors were matched to the decoys’ global relevance, and that decoys would not uniquely benefit from positive carry-over of attention from trial to trial, so that baseline and decoy distractors only differed in whether they were an anticipated target (future relevance). In contrast to Experiment 1, we now saw no reliable evidence of a difference between decoy and baseline conditions in either Target 1 (*d*=.13) or Distractor 1 weights (*d*=-.22; Figure 3). This suggests that the decoy results in Experiment 1 may have reflected the global or past relevance of the decoy colour, rather than its future relevance.

**Figure 3.**
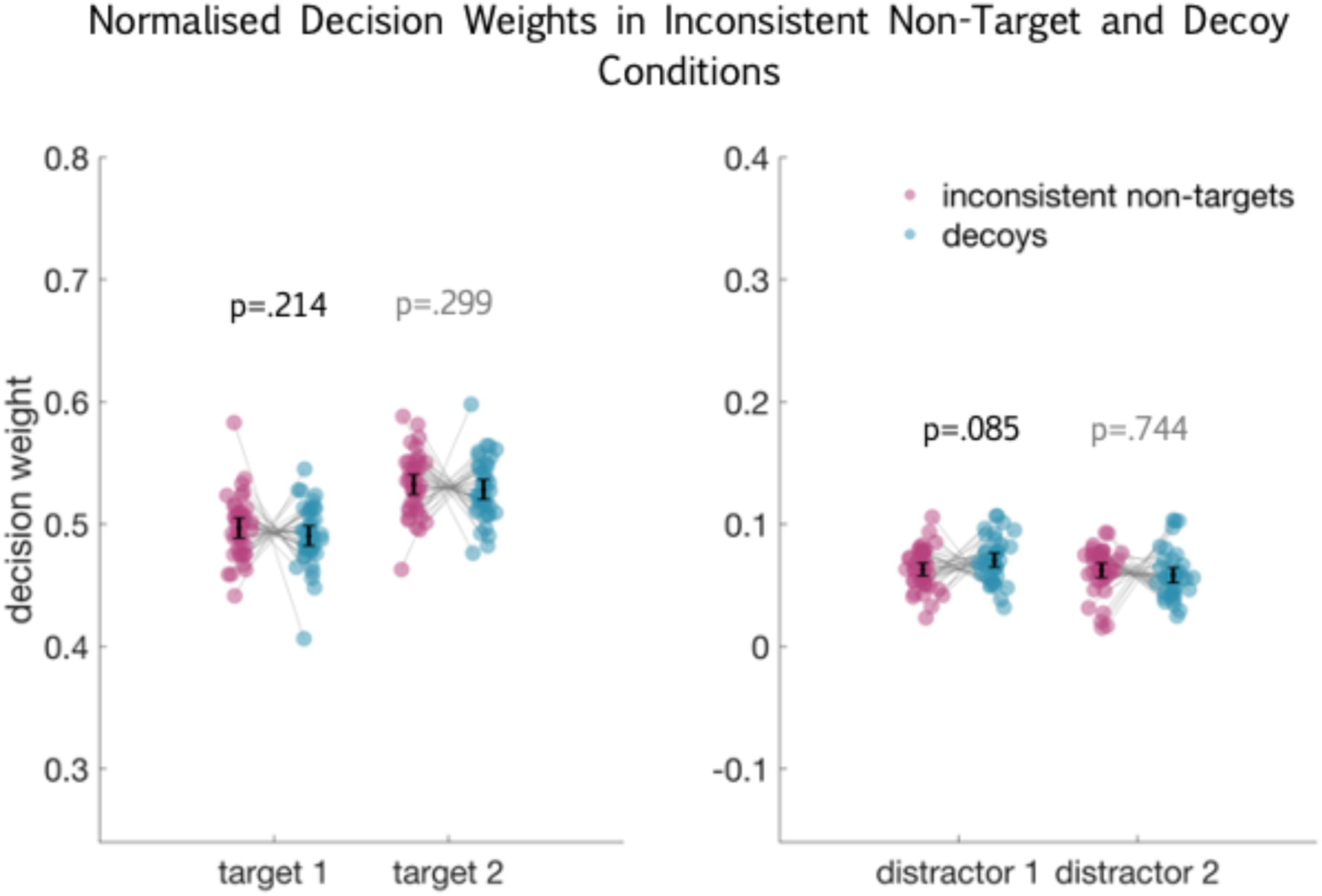
Normalised decision weights for target and distractor dot clouds in Experiment 2, shown separately for inconsistent non-target baseline (pink) and decoy (blue) conditions. Black bars indicate the 95% confidence interval. P-values are derived from one-sided paired t-tests and are uncorrected for multiple comparisons. Between-condition comparisons for Target 2 and Distractor 2 are in grey. There were no statistically significant differences between conditions.

On the other hand, failing to detect a decoy effect in Experiment 2 does not completely rule out the possibility that future-relevant information captures attention. Another possible explanation is that we failed to detect awareness of future-relevant task information in Experiment 2 simply because participants could no longer direct their attention to each display as it appeared. Although target weights were reliably different to zero, many participants spontaneously reported that they struggled to redirect attention to a new pair of target colours. Target weights were indeed lower in Experiment 2 relative to those in Experiment 1 (p=.038; Figure 4), based on an age-matched sub-sample (n=23, age=25.6±4.8 years, 13 female, 10 male). Participants could have compensated for the pressure to quickly switch focus between trials by taking a more retroactive approach to the task, relying on the stimuli to engage cognitive control rather than proactively maintaining the targets in mind (Braver, 2012). In a proactive mode, participants would adjust their attentional sets in advance so that only the task-relevant stimuli were encoded and maintained. In a retroactive mode, by contrast, participants would encode both relevant and irrelevant stimuli, and later, when presented with a response display, try to retrieve just the relevant stimuli. The latter might be a low-cost strategy (Braver, 2012), which could make it a natural choice when the participants were challenged by trial-to-trial switching demands. Under this framework, a more proactive approach in Experiment 1 could have enabled participants to quickly enhance anticipated targets and suppress consistent non-target distractors (in line with high performance in the baseline condition), but could fail when conflicting information was unexpected (in line with lower performance in the decoy condition). Thus, a proactive cognitive control strategy in Experiment 1 could produce low awareness of baseline distractors and high awareness of decoys, whereas a reactive control strategy in Experiment 2 could produce a stable, higher level of distraction across both conditions. If proactive and reactive control strategies produced the different results of Experiments 1 and 2, finding no decoy effect in Experiment 2 could still be consistent with a true decoy effect in Experiment 1.

**Figure 4.**
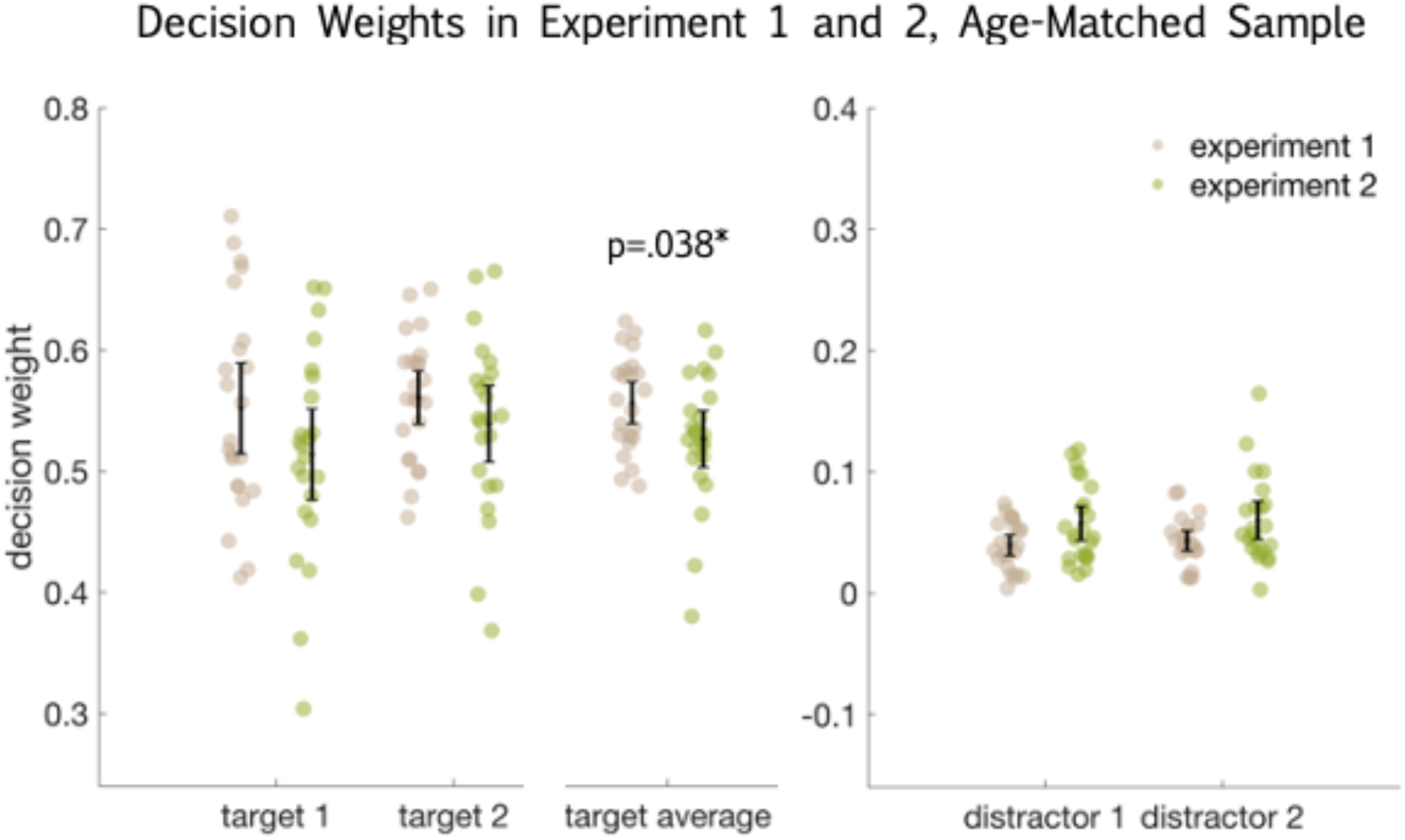
Average baseline decision weights for Targets 1 and 2 in Experiment 1 (grey) and an age-matched sub-sample of Experiment 2 (green). Decision weights are raw, not normalised, as this comparison is between samples. Black bars indicate the 95% confidence interval. The p-value for a two-sided, independent-samples t-test is shown below the average target weights (far right). We show baseline decision weights for target and distractor dot clouds in Experiment 1 (as in Figure 2) and the Experiment 2 subsample for completeness.

In addition, although there was no statistical decoy effect in Experiment 2 (despite a sample nearly twice that of Experiment 1) numerical trends were in the predicted direction. Within the sample, target weights tended to decrease and Distractor 1 weights tended to increase on decoy trials relative to baseline. Small and inconsistent changes do not support the conclusion that decoys are uniquely distracting, but neither do they rule it out. Given the ambiguity inherent in null effects, it is especially important that we consider how robust they are.

Considering these two limitations, we designed a third experiment to test the robustness of these findings, using a simplified task to promote proactive cognitive control.

## Experiment 3

So far, we have seen that future-relevant information may draw attention away from currently relevant information (Experiment 1), but that this bias toward future-relevant information is unreliable when we account for the distractor’s recent and global relevance (Experiment 2). However, Experiment 2 was difficult, with many participants spontaneously reporting that they struggled to switch focus between cues from trial to trial. We were also wary of over-interpreting the null finding in Experiment 2 without ensuring that we could replicate it. Therefore, in Experiment 3, we aimed to test the robustness of the finding from Experiment 2, using a simplified task and controlled trial sequence.

### Methods

#### Participants

Following the previous experiments, we conducted a power analysis to estimate an appropriate sample size. We maintained the same minimum effect of interest as in Experiment 2, again reasoning that distractor conditions separated by one aspect of distractor relevance should show smaller differences compared to the distinct distractor conditions in Experiment 1. However, as this experiment was designed in part to test the robustness of Experiment 2, and because testing online increased our pool of potential participants, we increased our target power to detect effects from .7 to .85. We again accounted for two planned within-subjects t-tests, alpha = .025 per test. G*Power suggested that approximately 75 participants would be sufficient to detect our minimum effect of interest.

We recruited an independent sample of 78 participants (age=28.6±9.7 years, 27 female, 51 male) for this study. Participants were recruited through the Prolific research recruitment portal (https://www.prolific.co/) and completed the task online. As in the previous experiments, the study was only advertised to participants who had previously reported that they were fluent in English, had normal or corrected to normal visual acuity, had normal colour vision, and were between the ages of 18 and 65. Additionally, participants were asked to join the experiment only on a desktop computer or laptop (not on a tablet or phone). All participants gave informed consent by clicking an online form. Ethical approval and payment remained the same as for Experiments 1 and 2, with no travel contribution offered now that participants could access the task online.

#### Task

As in Experiment 2, we used the same colours equally as targets and distractors (inconsistent non-targets), so that participants could not benefit from a global tendency to up-weight target colours. All methods were the same as in Experiment 2, with the following changes.

In Experiment 2, cues appeared in a random sequence, so that a decoy on the current trial was not consistently the second target on the previous trial. However, all four colours appeared on each trial, meaning that attentional biases toward each colour could carry over from trial to trial, albeit in positive (n and n-1 have the same targets), ambiguous (n and n-1 share only one target), and negative (n and n-1 have different targets) ways. For this experiment, we selected eight colours with hues 45° apart. We divided the eight colours into two sets, with hues 90° apart within each set (Figure 1B). We chose maximally distinct hues to ensure that, even if they were slightly altered by online participants’ computer monitors, they would be easily discriminated. We interleaved the two colour sets to create a train of four trials. Each train comprised (1) a trial from Set 1, for example, “orange, then blue”; (2) a trial from Set 2, for example, “purple, then pink”; (3) the inverse of Trial 1 (orange and blue as distractors); and (4) the inverse of Trial 2. We also increased the cue duration from 500 to 1000 ms to further support participants to prepare their attentional set on each trial.

We did not point out the trial sequence to participants, but they could learn it, explicitly or implicitly. Each block consisted of a single train, so that the exact combination of target and distractor colours repeated every five trials. Thus, we reduced the difficulty inherent in random cueing while making sure that the same colours were targets and distractors in both conditions.

Participants completed six blocks. Set 1 and Set 2 colours each formed six colour pairs (four colours combined without repetition; orange and blue, orange and pink, and so on), and each block contained a target colour pair from each set, so that every individual completed one block with each colour pair as the target colours. For a given block, we selected target colour pairs from each set to maximally vary within subjects what pairs were combined across sets. For example, if in Block 1 the Set 1 target colours were orange-green, and Set 2 target colours were dark blue-yellow, orange and dark blue would not be selected together as the Target 1 colours in a subsequent block, and the same for green and yellow. Combinations across colour sets were further randomised across participants.

These changes were designed to have two effects. First, we intended the extra preparation time and predictable trial sequence to simplify the task (Altmann, 2004; Longman et al., 2017; Meiran et al., 2000; Vandierendonck et al., 2010). Second, we intended the interleaved trials from distinct colour sets to further reduce the possibility of attentional carry-over, and the difficulty associated with conflicts in stimulus relevance from trial-to-trial (Gilbert & Shallice, 2002; Waszak et al., 2003). Relative to Experiment 1 these features again acted to isolate the effect of future relevance from possible effects of recent or global relevance. As before, we predicted that if momentary attention is affected by keeping track or of planning for a future attentional set, information relevant to a future task part (the decoy) would attract attention and disrupt performance, relative to the baseline distractors.

#### Stimuli

We generated random dot stimuli using bespoke JavaScript code and the jsPsych library (de Leeuw, 2015). We matched the dot cloud parameters (number, coherence, etc.) to Experiments 1 and 2.

For the response screen, we adapted the response dial to be compatible with jsPsych, and to be easy to see and manipulate with a mouse or touchpad. We used a grey circular dial with a white pointer (Figure 1, right). Feedback was presented as a light grey pointer overlaid on the white response pointer.

Participants moved the pointer with their mouse or touchpad, either by dragging the pointer or by clicking directly to where on the circle it should go. We reduced the maximum response time from 6000 ms to 3000 ms, as response times greater than this in Experiments 1 and 2 were rare.

Stimuli were presented on participants’ personal computers. We estimated each participant’s refresh rate by engaging a browser interface method for updating content on each screen “repaint” (typically the same as a screen refresh; see https://developer.mozilla.org/en-US/docs/Web/API/window/requestAnimationFrame), taking a timestamp on each repaint, and extracting the average time between repaints over five seconds. We also asked participants to match the size of an on-screen box to the size of a credit card. This gave us the pixel dimensions of an object with known true size, allowing us to estimate their screen resolution and match stimulus sizes across participants, devices and experiments. We adjusted the distance that each dot travelled on each frame to match speed across refresh rates and screen dimensions. As in the previous experiments, dot clouds covered a 14 cm circular area, so that the stimuli subtended 15.94 degrees of visual angle at a viewing distance of 50 cm. We asked participants to position themselves 50cm from the screen, but as we were unable to verify that they did so, the effective visual angle of the stimuli could have differed between participants and experiments. Due to participants accessing the study remotely, we also could no longer test how their screen distorted the colours. This meant that, although we set all colour values to 75% saturation and 50% luminance, true saturation and luminance may have varied across colours. However, we presented all colour pairs to each participant and conducted all primary analyses within-subjects, so that between-subject variability in displays would minimally influence the results.

We tested all stimulus presentation scripts locally and through JATOS (Lange et al., 2015), a web application that allows researchers to interact with a web server (for example, generate a study link and download data) through a graphical interface.

#### Procedure

We asked participants to sit 50 cm away from the screen, and to ensure that their screen faced them directly before beginning the task. We also presented a labelled image of the two colour sets (Figure 1B) prior to training, to remove any ambiguity about which cue word indicated which colour.

In Experiments 1 and 2, participants completed three training blocks of 24 trials each, with the cue duration beginning at 1000 ms in blocks one (single-epoch) and two (dual epoch) and reducing to the core session’s 500 ms in training block three. In the current experiment, the cue duration remained at 1000 ms throughout training and core sessions. Consequently, the third training block was not necessary to introduce a new cue duration. We therefore reduced the number of training blocks from three to two, but maintained the total number of training trials by increasing the number of trials per block from 24 to 32.

##### Data quality

Comparisons of web- and lab-based data quality have demonstrated that online platforms can be appropriate for cognitive psychology experiments (Germine et al., 2012), and that stimulus presentation times and participant compliance can compare positively with lab studies (using jsPsych and Prolific: Anwyl-Irvine et al., 2021; Peer et al., 2017). However, since we were unable to discuss the task with participants or adaptively respond to any issues they experienced, we implemented some additional measures to encourage and track data quality. Participants were encouraged to re-read the instructions for each training block. We could not offer individualised explanations or answer questions. Instead, we introduced additional tests to filter out participants who misunderstood the task or did not seriously attempt it. First, we included a catch question in the pre-experiment questionnaire (“What year is it?”), which blocked participants from continuing if they answered incorrectly. Next, we used errors on the single-epoch training block to identify underperforming participants. Whereas an error on dual-epoch trials could reflect bias toward one target over the other, errors on single-epoch trials showed that the participant had not accurately identified or perceived the target, making these errors a fair indicator of whether they understood and seriously attempted the task. Participants whose median absolute error in the first training block exceeded 45° (that is, most responses fell outside the quadrant for the correct response) were prevented from continuing and paid for their time. Lastly, participants who did not respond, or who responded with the default response dial position, for three sequential trials were shown a warning screen prompting them to respond within the time limit. Participants who received this warning three times, either during training or during the core session, were prevented from continuing and paid for their time. These measures allowed us to reject data from participants who responded with minimal effort, and also ensured that participants who were struggling did not continue to struggle through the intensive experiment session.

### Results

#### Baseline decision weights

Eleven participants were excluded for high errors, leaving 67 participants in the final sample (age = 27.67±8.93, 24 female, 43 male; 80% power to detect our effects of interest). All remaining subjects’ baseline target weights exceeded zero, relative to their individual permutation-based null distribution. Baseline target decision weights across the group were also reliably different from zero (p<.001). Baseline distractor decision weights were also reliably different to zero in Epoch 1 (p = .002), but not in Epoch 2 (p = .146).

#### Decoy effect on decision weights

In contrast to our prediction, and in line with Experiment 2, we observed no reliable change in Target 1 (*d*=.15) or Distractor 1 weights (*d*=.18) in the decoy condition (when Distractor 1 was the anticipated Target 2), compared with the baseline distractors (Figure 5). This was true despite low mean absolute errors (30.37°±6.56, compared with 32.67°±6.32 in Experiment 1). This replicates the null findings of Experiment 2 with an independent sample and an easier task with predictable cues. Together, the two experiments provide no evidence for the hypothesis that attention is directed toward task features that will shortly become relevant.

**Figure 5.**
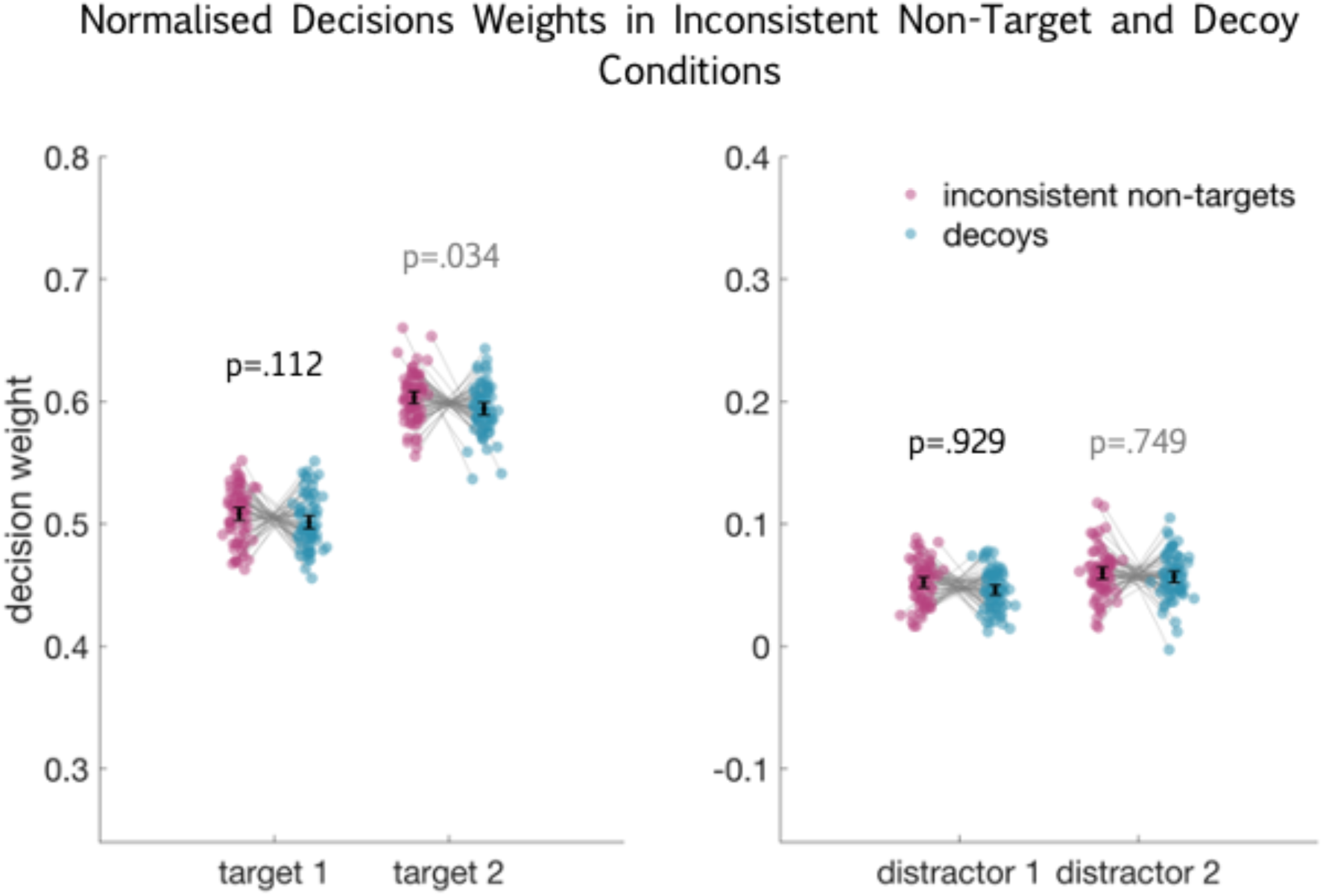
Normalised decision weights for target and distractor dot clouds in Experiment 3, shown separately for inconsistent non-target baseline (pink) and decoy (blue) conditions. Black bars indicate the 95% confidence interval. P-values are derived from one-sided paired t-tests and are reported without correction for multiple comparisons. The critical alpha for inconsistent non-target vs decoy conditions was .025, to account for planned comparisons of Target 1 and Distractor 1. We show outcomes for Target 2 and Distractor 2 (in grey) for completeness. There were no statistically significant differences between conditions.

In light of these findings, our new hypothesis was that future-relevant information does not capture attention beyond what we might expect from its relevance in the broader context of the task. Therefore, we designed a final experiment to (a) replicate the attentional capture by a decoy, relative to a consistent non-target, that we observed in Experiment 1; (b) directly test whether people indeed upweight globally relevant information (inconsistent non-targets and decoys) over consistent non-targets (as opposed to controlling for global and recent relevance in Experiments 2 and 3); and (c) directly compare the impact of global or recent relevance and future relevance within a sample.

## Experiment 4

Experiment 4 consisted of three conditions: a baseline condition with consistent non-target distractors (cNT), a baseline condition with inconsistent non-target distractors (colours drawn from the same set as target colours; iNT), and a decoy condition. We simplified the task further, relative to Experiment 3, by restricting the target/inconsistent non-target colour set to four colours, and using two (rather than four) cues in each block. As in all previous experiments, we presented every combination of colours to each participant, and randomised block sequence across participants.

In line with our new hypothesis that the broad experimental relevance of each colour determines attentional weighting, with little or no additional contribution of a colour’s imminent relevance later on that trial, we now predicted that the iNT condition would elicit lower average target weights, and higher average distractor weights, than the cNT condition. Based on Experiment 1, we predicted that decision weights in the decoy condition would also reflect poorer selectivity (lower target weights and higher distractor weights) compared with the cNT condition. Following Experiments 2 and 3, we now expected that Epoch 1 weights in the decoy condition would not differ from the iNT condition. This set of results would demonstrate that momentary attentional sets are sensitive to the global relevance of distractors, but do not suffer additional interference from information that will become relevant later in the same trial.

### Methods

#### Participants

Following Experiment 3, we set a target sample size of 75 participants. We recruited an independent sample of 76 participants (age=28.9±9.1 years, 25 female, 50 male, 1 unreported). Participants were recruited through Prolific and completed the task online. All participants gave informed consent by clicking an online form. Ethical approval and payment rate remained the same as for Experiments 1-3, with no travel contribution following Experiment 3. For this experiment, training was offered in a separate 15-minute session, for which participants received an additional £1.50.

#### Task

The task remained the same as in Experiment 3, with the following changes. We selected six colours, with hues 60° apart. For each participant, we assigned a pair of colours to be consistent non-target colours. The remaining four colours formed the target colour set. For each of six blocks, we assigned these four colours to two colour pairs. We balanced colour pairings across blocks so that each colour was paired at least once with the remaining three colours. On each trial, one colour pair was the cued target pair, and the other pair were the distractors. Colours within a pair could appear in any order when they served as distractors, but were always cued in a fixed order when they served as targets. This minimised the number of cue variations per block (set it equal to two), to reduce overall task difficulty. Trials in which the distractor pair were drawn from the held out colours formed a cNT baseline condition (similar to the baseline in Experiment 1). Trials in which the distractor pair were drawn from the target colour set formed an iNT baseline condition (similar to the baseline in Experiments 2 and 3). Decoy trials, in which the first distractor was also the anticipated target colour, made up a third condition. We believed that the iNT condition was a fairer baseline by which to isolate whether people were influenced by future relevant information, as both future relevant and inconsistent non-target colours were globally relevant for the block. Because of this, we always used inconsistent non-target colours as the second distractor in decoy trials.

Conditions and cues were presented randomly within each block. As the task now included three conditions, we reduced the length of each of the six core task blocks from 64 trials to 60 trials, so that we could include equal numbers of trials in each condition.

#### Stimuli

Screen resolution checks and stimuli were as described in Experiment 3, with the exception of the reduced colour set.

#### Procedure

The procedure was the same as in Experiment 3, with one added data quality assessment, described below.

##### Data quality

In Experiment 3, participants who passed quality check criteria were free to continue immediately to the core session. In Experiment 4, participants were additionally screened for technical issues (such as lags in the presentation) or personal issues (such as misunderstanding the task, finding it difficult, or being distracted), which they reported at the end of the training session. Participants who reported issues with the task were contacted to clarify the issue before being given the link to the core session. This was done to prevent people from continuing to the core session before they fully understood the task, and so expending substantial time and energy to give unusable data. The time between training and core sessions ranged from one hour to one week. Because of the delay between training and core sessions, the core session began with a full repetition of the instructions and three dual-epoch practice trials.

#### Analyses

As for all previous experiments, we controlled error rate family-wise, separately for baseline against zero, and for between-condition comparisons. We predicted that Target 1 and Distractor 1 weights would not change in the presence of a decoy distractor relative to the iNT baseline. The critical alpha for the decoy effect (iNT vs decoy conditions) was .025 and t-tests were two-sided. To test the impact of global relevance, we also contrasted decision weights between iNT and cNT conditions. Here, we expected that target decision weights would decrease and distractor decision weights would increase in both epochs, in the iNT condition relative to the cNT baseline. Thus, we set alpha at .013 for four planned, one-sided, comparisons. We did the same for decoy and cNT conditions, again predicting that target weights would decrease and distractor weights increase in both epochs, in the presence of globally relevant distractors.

### Results

#### Baseline decision weights

Five participants were excluded for high errors. One further participant’s cNT baseline target weights did not exceed the 95^th^ percentile in their permutation-based null distribution, meaning that their accuracy for at least one target could not be distinguished from chance. This participant was also excluded from subsequent analyses, leaving 70 participants in the final sample (age = 28.77±9.20 years, 22 female, 47 male, 1 unreported; 82% power to detect our effects of interest). Across the group, target and distractor weights in the cNT baseline condition were reliably above zero (all p<.001).

#### Decoy effect on decision weights

In line with Experiment 1, the decoy condition showed evidence of increased distraction (lower target weights, higher distractor weights) relative to the cNT baseline. However, as now predicted, and consistent with Experiments 2 and 3, decoy weights were again not reliably different to the iNT baseline (Table 1; Figure 6).

**Figure 6.**
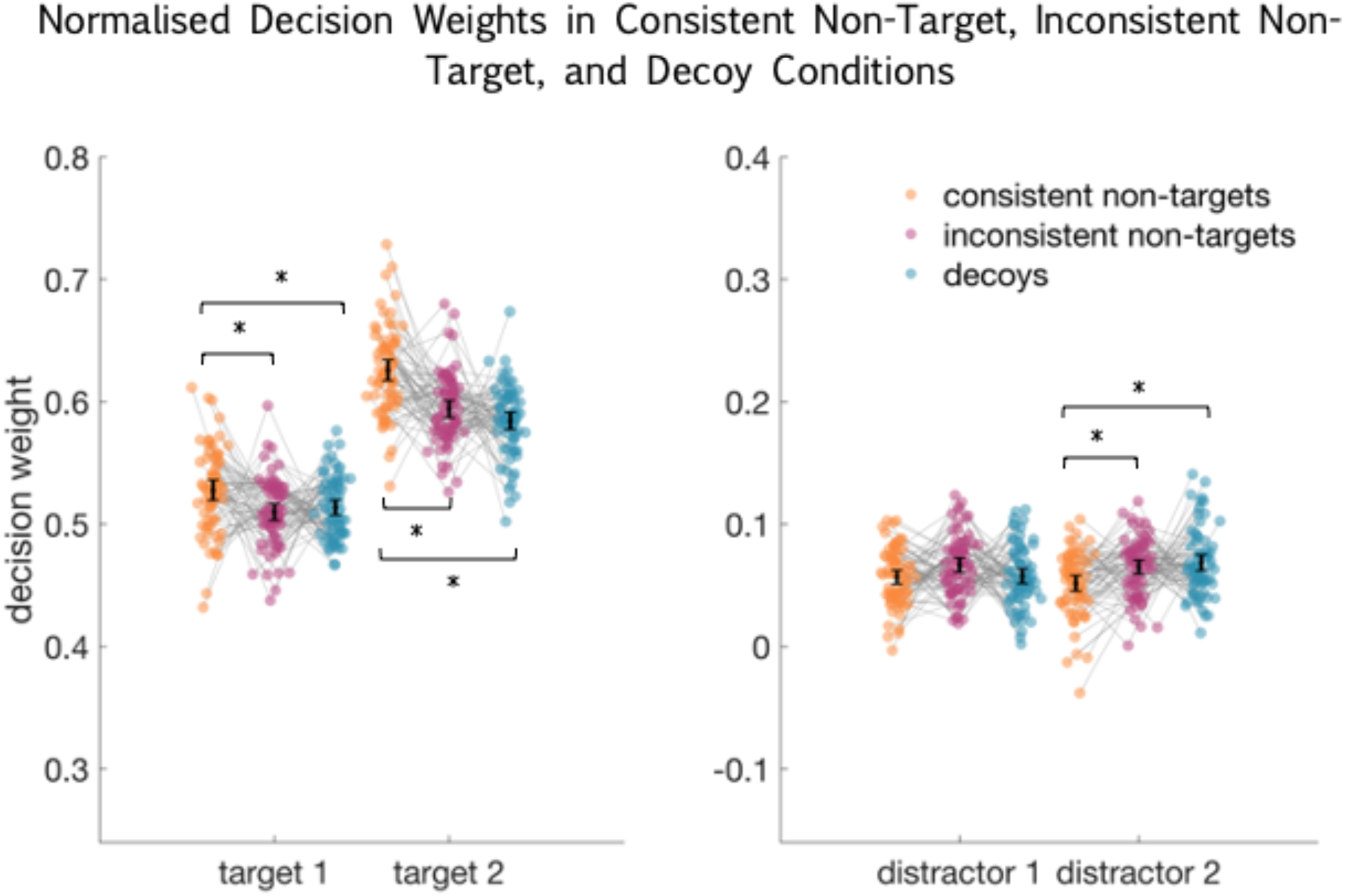
Normalised decision weights for target and distractor dot clouds for Experiment 4, shown separately for consistent non-target baseline (orange), inconsistent non-target baseline (pink), and decoy (blue) conditions. Black bars indicate the 95% confidence interval. Statistically significant condition differences are marked with an asterisk.

**Table 1.** Statistics for condition differences in decision weights, n=70

| comparison | Student's *t* | *p* | critical alpha | Cohen's *d* |
| --- | --- | --- | --- | --- |
| T1 cNT > T1 decoy | 2.43 | .009 | .013 | 0.29 |
| T2 cNT > T2 decoy | 6.22 | <.001 | .013 | .74 |
| D1 cNT < D1 decoy | -.17 | .431 | .013 | -.02 |
| D2 cNT < D2 decoy | -2.94 | .002 | .013 | -.35 |
| T1 iNT ≠ T1 decoy | -.63 | .533 | .025 | -.07 |
| T2 iNT ≠ T2 decoy | 1.49 | .141 | n/a | .18 |
| D1 iNT ≠ D1 decoy | 1.63 | .108 | .025 | .19 |
| D2 iNT ≠ D2 decoy | -.59 | .555 | n/a | -.07 |
| T1 cNT > T1 iNT | 2.69 | .004 | .013 | .32 |
| T2 cNT > T2 iNT | 4.80 | <.001 | .013 | .57 |
| D1 cNT < D1 iNT | -2.13 | .018 | .013 | -.25 |
| D2 cNT < D2 iNT | -3.07 | .002 | .013 | -.37 |

*Note. T2 and D2 decision weight comparisons for decoy and iNT conditions were not planned, but are shown for completeness.*

This experiment also directly confirmed that target weights in the iNT condition were lower, and distractor weights higher, relative to the cNT baseline. The finding was statistically significant in three out of four comparisons, across two epochs. This supports the suggestion that attention is captured both by what is immediately relevant and by what has been relevant (recently and/or globally). We did not find evidence to suggest that attention is additionally captured by information that will become relevant imminently.

#### Contribution of trial-to-trial attentional set vs global relevance

Through the previous experiments, we found that distractors substantially captured attention when they were future-relevant, recently relevant, and globally relevant (Exp 1). This effect disappeared when we controlled for recent and global relevance (Exps 2 and 3). The current experiment allows us to further separate these influences.

First, reliable differences between iNT and cNT conditions indicate that globally relevant distractors reliably capture attention, relative to distractors that are never relevant. The global relevance effect persists even when what is relevant changes pseudo-randomly from trial to trial (with equal opportunities for recent relevance to help or hinder current focus), suggesting that the global relevance effect is unlikely to rely on trial-to-trial carry-over of attention.

However, it is possible that trial-to-trial carry-over effects are asymmetrical, making it difficult to rule them out completely in the current experiment, or in Experiment 1. We can instead quantify their effect by extracting the decision weights for cNT and iNT conditions, separately for whether targets repeat (“stay”) or change (“switch”). For both cNT and iNT conditions, target weights on the current trial could benefit from a repetition (“stay”), or be disadvantaged by a change (“switch”), as the same colours could be enhanced across trials. Distractor weights could also benefit from repetition, as the same colours could be suppressed across trials. However, the potential for distractor carry-over in switch trials would be different for cNT and iNT conditions. For the cNT condition, distractors could change or repeat in switch trials, but current trial distractors were never previous-trial targets; they were always consistent non-targets. Thus, carry-over from the previous trial could aid in suppressing the distractors (where two cNT trials occurred in sequence), but should not enhance the distractors. For the iNT condition, however, switching targets always co-occurred with the previous trial’s target colours becoming the current trial’s distractors. If attention to targets on the previous trial drove awareness of them on the current trial, we expected that target weights would decrease in both cNT and iNT conditions in switch trials, relative to stay trials, but that distractor weights would increase only in iNT switch trials relative to iNT stay trials. We excluded the decoy condition from this comparison, because decoy “stay” trials conflated potentially beneficial carry-over (repeating targets) with potentially harmful carry-over (trial n-1 Target 2 becomes trial n Distractor 1).

To understand how recent and global relevance drive attentional capture, we ran an exploratory repeated-measures analysis of variance on the decision weights, with global relevance (cNT vs iNT), target carry-over (stay vs switch), and dot cloud (T1, T2, D1, D2) as within-subject factors. The interaction between all three predictors was statistically significant (F_1,67_=7.90; p=.006), suggesting that global relevance and switch/stay have some effect on the decision weights, but that this effect differed between the dot clouds (Figure 7).

**Figure 7.**
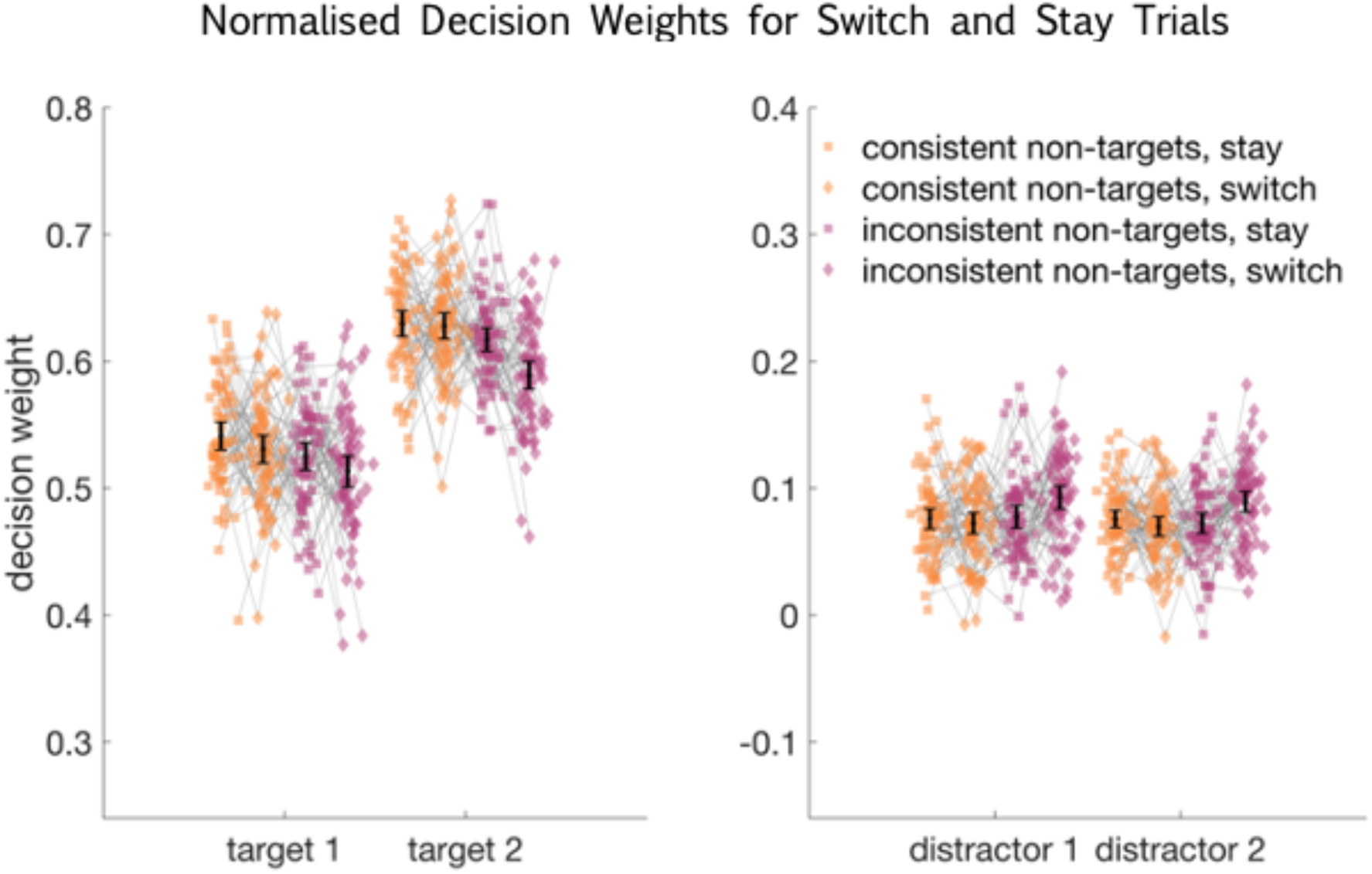
Normalised decision weights for target and distractor dot clouds for Experiment 4, shown separately for consistent non-target (orange) and inconsistent non-target (pink) stay and switch conditions (square and diamond markers respectively). Black bars indicate the 95% confidence interval.

Recall that while targets could switch or stay in cNT and iNT conditions, distractors on iNT switch trials were also always the previous target colour. This was never the case for distractors on cNT switch trials. If trial-to-trial carry-over of attentional set primarily influenced attention to targets, we could expect to see different target decision weights on stay and switch trials, with no reliable effect among distractors. If trial-to-trial carry-over of attentional set also increased attention to distractors that had recently been targets, we would expect to see high distractor weights on iNT switch trials, relative to all other trials.

So, we conducted follow-up analyses, now considering target and distractor weights separately. First, we re-ran the previous ANOVA, but with dot cloud divided into two factors: target status (targets vs distractors) and epoch (epoch 1 vs epoch 2). We saw a significant main effect of epoch (F_1,67_=42.45; p<.001) and a significant interaction between epoch and target status (F_1,67_=52.83; p<.001). However, epoch was not reliably associated with global or recent relevance, in direct, three-way, or four-way interactions. Consequently, we averaged decision weights across epochs to simplify the analysis of relevance effects. We then conducted two ANOVAs, one for target decision weights, and another for distractor decision weights, with global and recent relevance as predictors. We found that target decision weights were additively influenced by whether the distractors were globally relevant (F_1,67_=25.39; p<.001), and by whether targets repeated or changed relative to the previous trial (F_1,67_=10.88; p=.002), with no reliable interaction (F=2.43; p=.124). For distractors, however, the influence of global relevance and target switch was best described by a significant interaction (F_1,67_=9.70; p=.003). A planned comparison of iNT switch trials to all other conditions revealed that distractor weights increased when they were targets on the previous trial (t_67_=3.89; p<.001). This highlights two different effects of recent target relevance: Target weights drop following a switch, regardless of what distractors appear, whereas distractor weights selectively increase when the previous targets become the current distractors. This highlights the complex role of global and recent history in driving attention toward stimuli that are repeatedly relevant.

## Discussion

Attending to what is immediately relevant is a useful tool for allocating our mental resources toward simple parts of a task. Conversely, attending to features that may shortly become relevant for subsequent task parts could help us shift our focus when we need to. Here, we conducted four experiments to probe whether our attention is captured by irrelevant information if it will shortly become relevant. In Experiment 1, we observed evidence of future-task awareness, when future-task relevant stimuli were also globally and recently relevant. However, in three further experiments, we showed that the effect in Experiment 1 can be accounted for by enhanced attention to distracting information that is broadly relevant in the overall context of the task and/or recently attended, with little or no additional effect of something becoming relevant imminently. That is, we are sensitive to the past and/or global relevance of currently-irrelevant information, but do not appear to give preference to information that is imminently relevant for a future task part. These results emphasise our capacity for both sensitivity to broad attentional relevance, and temporally precise focus, as we move through parts of a task.

Many lines of inquiry have shown that we pay attention to what has been relevant in the past, both globally, over an extended period of time, as well as recently, in the immediately preceding trial. Repeating stimuli prime our perception of their colour, motion, and identity (Ellis et al., 1987; Kristjánsson & Campana, 2010; Maljkovic & Nakayama, 1994), and inter-trial sequence effects can even lead to us misperceive objects in a series as having similar features (Fischer & Whitney, 2014; Maloney et al., 2005). Priming is a perceptual effect, meaning that we perceive current targets more quickly and accurately when they match previous targets. However, rapid and precise perception could in turn drive rapid and precise responses to those stimuli. The current study is consistent with this idea, showing that we select relevant target colours more accurately when cues repeat over trials (Exp 4).

However, a purely perceptual bias is unlikely to drive the consistent preference for recurring targets that we report here. Each trial in this multi-part design included four distinct colours and pseudo-randomly varying motion directions. This means that adjacent perceptual features (for example, trial n-1 Target 2 and trial n Target 1 colour) only reliably repeated in the decoy condition of Experiment 1 and in decoy “stay” trials of Experiment 4. Any serial perceptual effects within the decoy condition of Experiment 4 were not sufficient to reliably increase attentional capture by distracting information, relative to baseline (decoy vs iNT, Figure 6).

Beyond perceptual bias, this study highlights the impact of previous task set on current target and distractor processing. Directing our attention towards new information requires that we disengage from our previous attentional set and reconfigure our attention to prioritise the new targets (Imburgio & Orr, 2021). Task-switching studies have repeatedly demonstrated that this process is non-trivial, with task-set inertia biasing our focus towards what was recently relevant, and imposed time limits making it difficult for us to assemble our new task set (Evans et al., 2015; Imburgio & Orr, 2021; Longman et al., 2017; Weiler et al., 2015). The present study offers an added level of specificity to previous studies of errors in task-switching, by separately tracking responses to targets and distractors. Here, we found that responses reflect less target information when targets switch. Curiously, responses did not correspondingly reflect more distractor information on target switch trials. Instead, distractors were only upweighted on switch trials, relative to stay trials, when they had been relevant on the previous trial.

We also observed a tendency to upweight colours that were sometimes targets (inconsistent non-targets), relative to colours that were never targets (consistent non-targets). This cannot be explained by colour salience, as we randomly assigned colours to non-target and inconsistent non-target colour sets for each participant. Instead, it could reflect reinforcement learning, as participants learned that attending to targets and suppressing distractors led to positive feedback at the end of the trial. This in turn could drive a policy of increasing attention to globally relevant colours (Botvinick, 2012; Sutton & Barto, 2018). In Experiment 4, the reinforced target colours could also be distractors (inconsistent non-targets), but the policy of preferring to respond to those colours would on average be beneficial. These four colours were more likely to be targets than distractors at a rate of 3:2, as they appeared as targets in all three conditions and distractors only in two. While reinforcement learning is often discussed in the context of associating stimuli with explicit actions, this study emphasises our capacity to associate stimuli with mental actions, such as “attend” or “ignore”.

Neural data from non-human primates offer a time-resolved interpretation for preferential responses to globally relevant distractors. Neurons in monkey prefrontal cortex commonly code information that is behaviourally relevant (Duncan, 2001; Erez et al., 2020; Erez & Duncan, 2015; Kadohisa et al., 2015; Stokes, 2011). However, these neurons initially code relevant and irrelevant information in multi-item displays, before giving priority to the relevant feature (Kadohisa et al., 2013). Interestingly, these data showed that neural coding of globally relevant distracting information (i.e., inconsistent non-targets) disappears less efficiently and completely, compared with coding of features that are never relevant (consistent non-targets).

We should acknowledge that our finding in Experiment 4, that any increased distraction by future-relevant colours can be explained by that colour’s previous use as a target, may not fully capture the effect we saw in Experiment 1. In Experiment 1, decoys reliably skewed the response away from the targets and toward the distractors, relative to trials on which the distractor was a consistently irrelevant colour. In Experiment 4, we created conditions to mirror Experiment 1, but retained design changes from Experiments 2 and 3 (written cues, target colours changing throughout the block). While it is difficult to completely rule out the possibility that decoy effects in Experiment 1 partly arose from the decoy’s future relevance, based on the near identical responses to trials containing decoys or inconsistent non-targets in Experiment 4, we believe that this is unlikely.

One limitation of this study comes from the choice of decoy distractor. We have argued that people do not appear to direct their attention toward future-relevant information as they move through parts of a task, based on a task in which we deliberately gave people access to task-relevant information outside the epoch in which it was relevant. But while the colour of the decoy was the future target colour, those dots contained no useful information about the direction of the future target dots. We might better represent the real world by making the information for all task parts available at the same time, and asking people to self-select the sequence of task parts. This self-directed aspect is present in matrix reasoning and block design tasks that are often used to probe fluid ability, and could be a primary reason that those tasks are so difficult (Duncan et al., 2017). However, introducing participant-driven task epochs presents some challenges for controlled research. We often rely on carefully constructed displays, and specific presentation times, to infer what people are thinking of at any moment. In the current task design, showing both target dot directions together would make it difficult to extract the temporal information we care about, that is, whether attending to each target in strict sequence leads to better performance. Possible ways forward would be to use eye-tracking, interactive stimuli, or time-resolved neural data with encoding models to gain precise information about what is being attended to at each point in time. Along with the current study, this could give us insight into how we are able to direct our mental resources efficiently toward our task goals as we interact with the dynamic world around us.

## Rights retention statement

For the purpose of open access, the author has applied a Creative Commons Attribution (CC BY) licence to any Author Accepted Manuscript version arising from this submission.

